# *In silico* simulations of erythrocyte aquaporins with quantitative *in vitro* validation

**DOI:** 10.1101/2020.05.02.074591

**Authors:** Ruth Chan, Michael Falato, Huiyun Liang, Liao Y. Chen

## Abstract

Modelling water and membrane lipids is an essential element in the computational research of biophysical/biochemical processes such as water transport across the cell membrane. In this study, we examined the accuracies of two popular water models, TIP3P and TIP4P, in the molecular dynamics simulations of erythrocyte aquaporins (AQP1 and AQP3). We modelled the erythrocyte membrane as an asymmetric lipid bilayer with appropriate lipid compositions of its inner and outer leaflet, in comparison with a symmetric lipid bilayer of a single lipid type. We computed the AQP1/3 permeabilities with the transition state theory with full correction for recrossing events. We also conducted cell swelling assays for water transport across the erythrocyte membrane. The experimental results agree with the TIP3P water-erythrocyte membrane model, in confirmation of the expected accuracy of the erythrocyte membrane model, the TIP3P water model, and the CHARMM parameters for water-protein interactions.

## INTRODUCTION

Transport of water across the cellular membrane is an essential biological process, which is facilitated by the water channel proteins, aquaporins (AQPs) ^1-8^. Naturally, the literature on AQPs is extremely rich and the research is currently active. See, *e.g., Refs.* ^9-31^ for some recent studies. Alongside experimental investigations, molecular dynamics (MD) simulations have been carried out to elucidate the molecular/atomistic insights that cannot be gained from experiments alone^32-39^. Now the copy numbers of the AQPs natively expressed in the human erythrocyte are known quantitatively^40^, can we make quantitatively accurate predictions of their transport characteristics from the all-atom MD simulations? Is it necessary to model the erythrocyte membrane with appropriate lipid compositions in its inner and outer leaflets^41-46^? In another word, what difference would it be if the membrane is simply modelled as a lipid bilayer of a single lipid type as it is in the current literature? Relatedly, which simple water model, TIP3P^47, 48^ or TIP4P/2005^49, 50^, gives a more accurate representation of the essential solvent, water, in the study of water channels? In this article, we answer these questions with the following *in silico* and *in vitro* investigations.

We conducted all-atom MD simulations of AQP1 and AQP3 that are natively expressed in the human erythrocyte. Each model system consists of a biologically functional unit of the channel proteins (AQP1/3 tetramer) embedded in a patch of erythrocyte (noted as RBC hereafter) membrane with appropriate lipid composition tabulated in Table I (illustrated in Fig. 1). We used both the TIP3P water model and the TIP4P/2005 water model in respective studies. TIP3P was found to be more accurate for water transport through AQPs than TIP4P/2005 even though the latter gives better estimate for the bulk water properties. This is understandable because water conduction through AQPs is largely determined by the protein-water interactions represented by the CHARMM force parameters that are optimized with the TIP3P water model. For comparison, we also conducted simulations of AQP1 and AQP3 embedded in a patch of phosphatidylethanolamine (POPE) bilayer using TIP3P. In quantitative differentiations of the all-atom MD simulations, we conducted *in vitro* experiments on human erythrocytes. Using light scattering off the erythrocyte mixtures in a stopped-flow device, we quantified the cellular transport process with known accuracy. The experimental data validate our *in silico* study of erythrocyte membrane model with TIP3P and CHARMM36 parameters. The experiments further illustrate that AQPs are not modulated by the erythrocyte conformational changes.

**Table I.** Lipid compositions of the model erythrocyte membrane (RBC).

|  | Cholesterol (CHL) | Phosphatidylcholine (POPC) | Phosphatidylethanolamine (POPE) | Phosphatidylserine (POPS) | Sphingomyelin (SSM) |
| --- | --- | --- | --- | --- | --- |
| Inner leaflet | 20% | 11% | 38% | 22% | 9% |
| Outer leaflet | 20% | 35% | 10% | 0% | 35% |

**Fig. 1.**
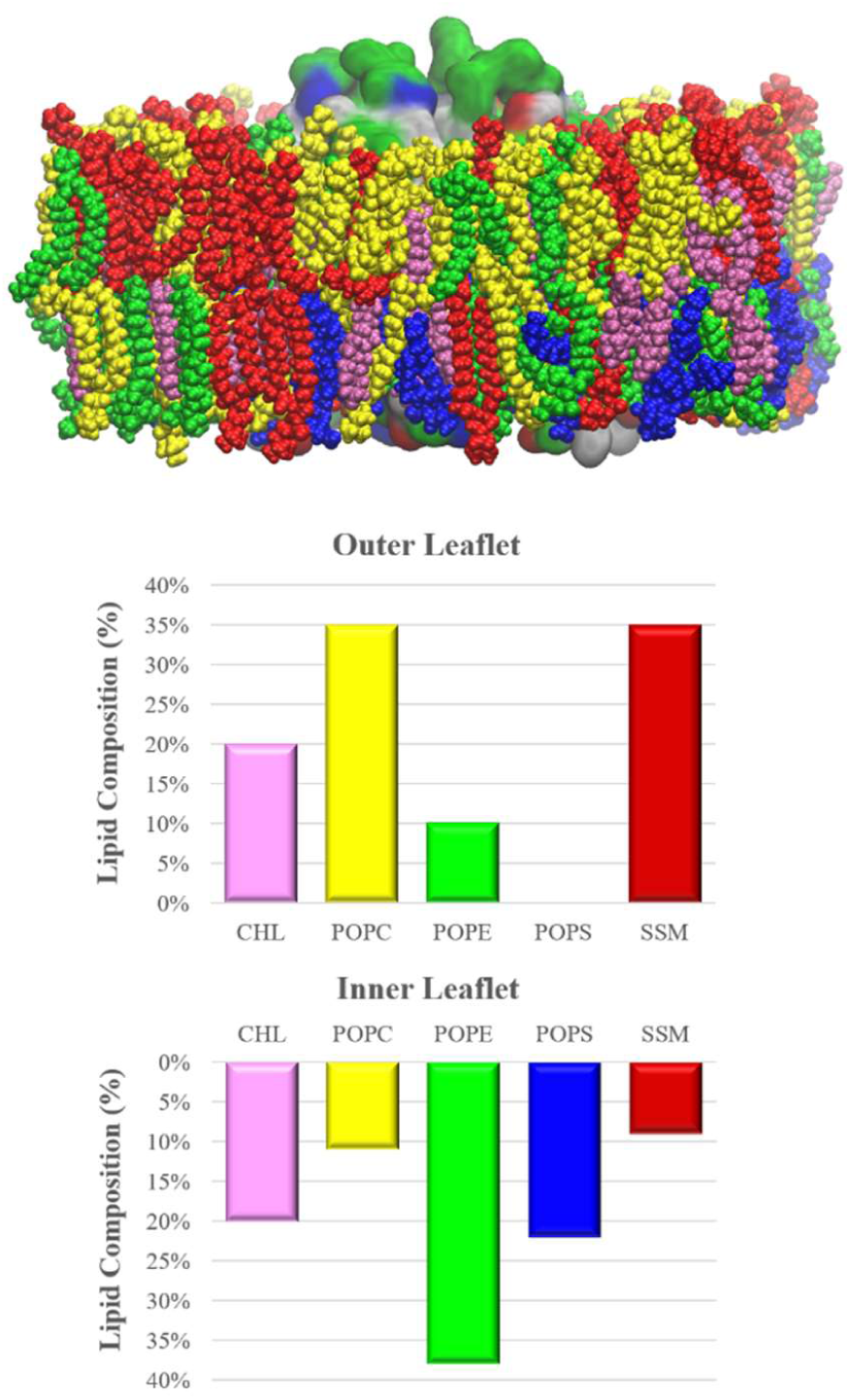
Model erythrocyte membrane. Lipids are shown as space-filling spheres colored by lipid types. Molar percentages and colors are shown in the bottom two panels. Four AQP1 monomers are shown in surface colored by residue types (hydrophilic, green; hydrophobic, white; negatively charged, red; positively charged, blue). All the molecular graphics of this article were rendered with VMD^51^.

## EXPERIMENTAL METHODS

### System setup and simulation parameters

Following the well-tested steps of the literature, we employed CHARMM-GUI^52^ to set up six all-atom model systems tabulated in Table II. In each system, the AQP1/3 tetramer was embedded in a patch of membrane. The AQP-membrane complex was sandwiched by two layers of TIP3P/TIP4P waters, each of which is approximately 35Å thick. The system was then neutralized and salinated with Na? and Cl^−^ ions to a salt concentration of 150 mM. A representative system is illustrated in Fig. 2. We employed NAMD 2.13^53^ as the MD engine. We used CHARMM36 parameters^54-56^ and the TIP3P or TIP4P/2005 parameters for inter- and intra-molecular interactions. After the initial equilibration steps, we conducted unbiased MD production runs (150 ns for each system) with constant pressure at 1.0 bar (Nose-Hoover barostat) and constant temperature at 298.15 K (Langevin thermostat). The Langevin damping coefficient was chosen to be 1/ps. The periodic boundary conditions were applied to all three dimensions. The particle mesh Ewald was used for the long-range electrostatic interactions (grid level: 128×128×128). The time step was 2.0 fs. The cut-off for long-range interactions was set to 10 Å with a switching distance of 9 Å. The last 100 ns of the trajectories were used in the computation of transport characteristics. (All the coordinates, parameters, and scripts necessary to reproduce the results of this study is available at https://doi.org/10.7910/.)

**Table II.**
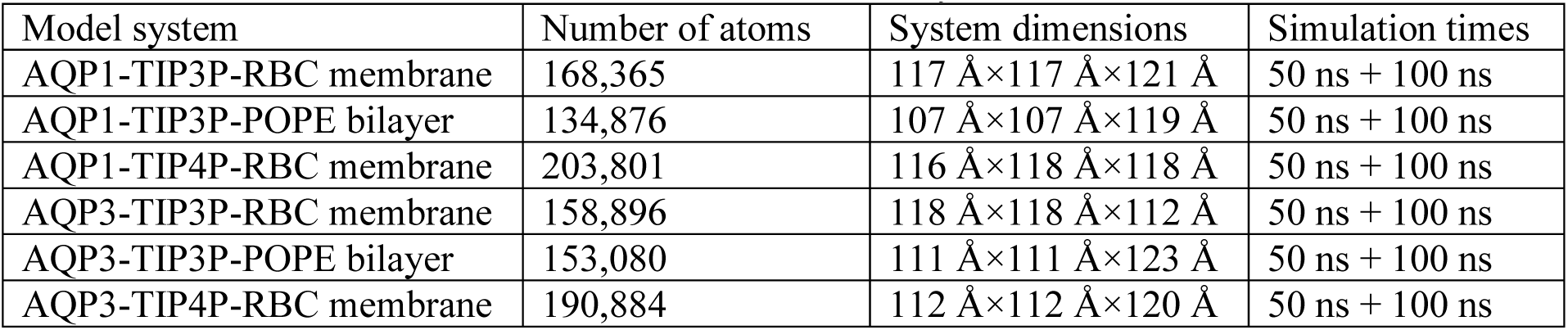
Simulation systems.

| Model system | Number of atoms | System dimensions | Simulation times |
| --- | --- | --- | --- |
| AQP1-TIP3P-RBC membrane | 168,365 | 117 Å×117 Å×121 Å | 50 ns + 100 ns |
| AQP1-TIP3P-POPE bilayer | 134,876 | 107 Å×107 Å×119 Å | 50 ns + 100 ns |
| AQP1-TIP4P-RBC membrane | 203,801 | 116 Å×118 Å×118 Å | 50 ns + 100 ns |
| AQP3-TIP3P-RBC membrane | 158,896 | 118 Å×118 Å×112 Å | 50 ns + 100 ns |
| AQP3-TIP3P-POPE bilayer | 153,080 | 111 Å×111 Å×123 Å | 50 ns + 100 ns |
| AQP3-TIP4P-RBC membrane | 190,884 | 112 Å×112 Å×120 Å | 50 ns + 100 ns |

**Fig. 2.**
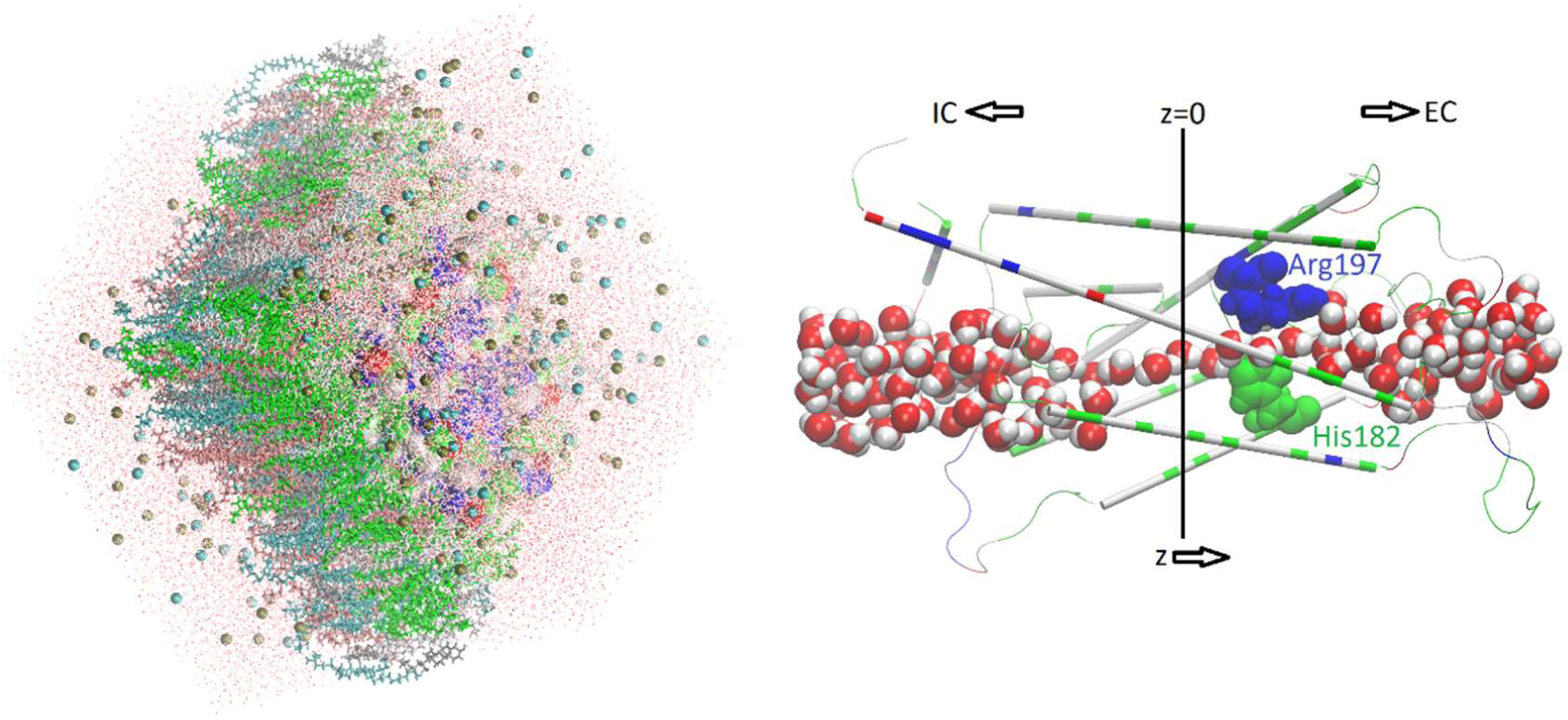
All-atom model system of AQP1 tetramer in erythrocyte membrane solvated with TIP3P waters. The full system is shown in the left panel where waters are in lines colored by atoms (O, red; H, white); NaCl ions are shown in spheres colored by atoms; protein in surface colored by residue types; and lipids in licorices colored by residue names. In the right panel, one monomer channel protein is shown in thin cartoons with the aromatic/arginine selectivity filter (ar/R sf) residues (Arg 197 and His 182) in large spheres, all colored by residue types. Water molecules inside and near the channel are shown in large spheres colored by atoms. The z-axis points from the intracellular side (z<-15Å) to the extracellular side (z>15Å).

### Transition state theory for water permeability

It has been clearly illustrated in the literature that an imbalance of osmolyte concentrations between the two sides of a membrane generates a difference/gradient in the chemical potential of water molecules. It is such a chemical potential gradient that drives the osmotic flux of water across the membrane (through AQPs) that is impermeable to the osmolyte solutes. Specifically, consider the extracellular (EC) and the intracellular (IC) sides of an erythrocyte and use *c*_*e*_ and *c*_*i*_, respectively, to denote the EC and the IC concentrations of the impermeable solutes. The difference in water chemical potential between the two sides^33^, *μ*_*e*_ − *μ*_*i*_ = (*p*_*e*_ − *p*_*i*_)*v*_*W*_ − *RT*(*c*_*e*_ − *c*_*i*_)*v*_*W*_. Here *p*_*e*_ − *p*_*i*_ is the difference in mechanic/hydraulic pressure between the two sides; *R* is the gas constant; *T* is the absolute temperature; and *v*_*W*_ is the molar volume of water. The IC-to-EC transition rate constant of water *k*_*ltoE*_ is related to the EC-to-IC rate constant *k*_*Etol*_: *k*_*ltoE*_ = *k*_*Etol*_exp [−(*μ*_*e*_ − *μ*_*i*_)/*RT*]. In the absence of a hydraulic pressure difference, *p*_*e*_ − *p*_*i*_ = 0, *k*_*ltoE*_ = *k*_*Etol*_exp [(*c*_*e*_ − *c*_*i*_)*v*_*W*_]. Under a hyperosmotic condition, *c*_*e*_ − *c*_*i*_ > 0, *k*_*ltoE*_ > *k*_*Etol*_. In this case, the outward chemical potential gradient induces a net efflux of water. The corresponding transition rate (transitions per unit time facilitated by one AQP channel) *r* = *k*_*ltoE*_(1/*v*_*W*_ − *c*_*i*_) − *k*_*Etol*_(1/*v*_*W*_ − *c*_*e*_). Note that 1/*v*_*W*_ is the concentration of water molecules in the absence of solutes. The presence of solutes reduces the water concentration. Furthermore, the validity of these formulas is limited to the dilute solution regime, namely, (*c*_*e*_ − *c*_*i*_)*v*_*W*_ ≪ 1, which is quantitatively accurate for osmolyte concentration in the sub Molar range. In this regime, the linear expansion in terms of (*c*_*e*_ − *c*_*i*_) is valid, which leads to the transition rate *r* = 2*k*_0_(*c*_*e*_ − *c*_*i*_). Therefore, the water flux through a single AQP channel (the volume of water flowing through a channel per unit time), *J* = *rv*_*W*_/*N*_*A*_ = 2*k*_0_ (*c*_*e*_ − *c*_*i*_)*v*_*W*_/*N*_*A*_. Correspondingly, the single-channel permeability *p*_*f*_ = *J*/(*c*_*e*_ − *c*_*i*_) = 2*k*_0_*v*_*W*_/*N*_*A*_. Here *k*_0_ is the rate constant *k*_0_ = *k*_*Etol*_ = *k*_*ltoE*_ at equilibrium (*c*_*e*_ = *c*_*i*_) and *N*_*A*_ is the Avogadro number.

However, the rate constant of IC-to-EC or EC-to-IC transition cannot be simply computed as the rate of water molecules crossing the dividing plane between the IC and the EC sides (z=0, Fig. 2) because the potential energy landscape of the water-protein-membrane system is very rugged^57^ and thus there are many recrossing events that need to be considered^58, 59^ in the application of the transition state theory (TST) to compute the transition rate constant^60^. Many of the events of a water molecule crossing the dividing plane actually end up recrossing the plane in the opposite direction and thus should be excluded from the counting of transitions between the IC and the EC sides. The IC-to-EC transitions consist of only the events of crossing the dividing plane in the positive z-direction and continuing onto the EC side without recrossing the dividing plane. Likewise, the EC-to-IC transitions. Therefore, TST gives 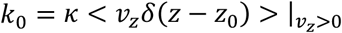 where *κ* is the correction factor and *z* = *z*_0_=0 (Fig. 2) is the dividing plane. The brackets indicate equilibrium statistical average of the velocity along the z-direction (or, equivalently, the negative z-direction for the EC-to-IC rate constant. Noting again that *k*_0_ = *k*_*Etol*_ = *k*_*ltoE*_ at equilibrium). Carrying out the equilibrium statistical average, we have 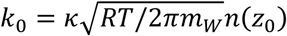 with *m*_*W*_ being the molar mass of water. The linear density of water at the dividing plane *n*(*z*_0_) can be readily evaluated with equilibrium sampling. The evaluation of the correction factor *κ* is computationally expensive. *κ* = 1 when the dividing plane *z* = *z*_0_ is the perfect transition state for a water molecule. If we had such a perfect transition state, once a molecule crosses the dividing plane along the z-direction, it will go on to the EC bulk. *Vice versa*, once a molecule crosses the dividing plane along the opposite direction, it will go on to the IC bulk. In our study of water transport through an AQP channel, there is practically not a perfect transition state and, as such, we did not try to locate it. Instead, we evaluated the correction factor *κ* in the following manner: For each event of a molecule crossing the dividing plane from the IC side (*z* < 0) to the EC side (*z* > 0), we followed its trajectory until it either recrosses the plane back to the IC side or moves out of the channel into the EC bulk (*z* > 15). For each event of a molecule crossing the plane along the opposite direction, we followed its trajectory until it either recrosses the plane back to the EC side or goes on to the IC bulk (*z* < −15). The correction factor *κ* is equal to the number of successful transport events (ending up in the EC/IC bulk) divided by the total number of events of crossing the dividing plane. We reached convergence of the statistics for the correction factor and the linear density. In this way, we computed the single-channel permeability of an AQP with known margin of error. It is gratifying to note that the final result of permeability so obtained is insensitive to the exact location of the dividing plane as long as *z*_0_ is inside the channel.

### Experimental procedure and analysis

The packed red blood cells from three anonymous healthy donors were purchased from the South Texas Blood and Tissue Center. Equal volumes of the three erythrocyte samples were mixed and washed three times and then suspended in phosphate buffered saline (PBS) to a total concentration of 4% hematocrit. In each experimental run, 5 µL of the 4% hematocrit suspension in PBS was rapidly mixed with an equal volume of PBS containing 180 mM sucrose using an Applied PhotoPhysics SX20 stopped-flow spectrometer. In such a mixture solution, the extracellular environment of the erythrocyte is hyperosmotic with a gradient of 90 mM which drives a water flux out of the cell resulting in cellular shrinkage. The intensity of light scattered at 90° was measured to monitor the erythrocyte shrinking process. The scattered light intensity is related to the varying cellular volume as follows:

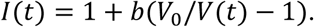

Here *t* is time. *b* is a parameter to account for the number of erythrocytes in the samples of a given set of experiments. *V*(*t*) is the intracellular volume of an erythrocyte with its initial value noted as *V*_0_. It follows the following dynamics equation:

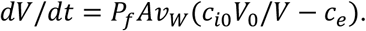

Here *P*_*f*_ is the osmotic permeability of the entire cell and *A* is the cellular surface area. *c*_*i*0_ is the initial intracellular concentration of impermeable solutes.

In an exponential fitting, we followed what is customary in the literature and assumed that the scattered light intensity to be an exponential function of time *I*(*t*) = *A* + *B* exp [−*k*_*fit*_*t*] that has three fitting parameters *A, B*, and *k*_*fit*_. We conducted least squares fit to obtain the rate *k*_*fit*_ which is assumed to be proportional to the cellular permeability. It has been pointed out in the literature that the exponential fitting for cellular permeability can be problematic^61^.

In contrast to the exponential fit, we numerically integrated the differential equation of the full dynamics for the time course of scattered light intensity *I(t)* for each set of values for the two fitting parameters *b* and *P*_*f*_*A*. All the other quantities involved (the water molar volume, the initial volume of the erythrocyte, the intracellular and extracellular concentrations of the impermeable solutes) were either known or measured at the time of experiment. We conducted least squares fit to obtain the mean and the standard error of the total cellular permeability *P*_*f*_*A* without any other assumptions.

## RESULTS AND DISCUSSION

### Bulk water diffusion

To confirm the literature on the bulk water diffusion constants, we conducted 100 ns MD simulations for a 100Å × 100Å × 100Å box of water represented as TIP3P^47, 48^ and TIP4P/2005^49, 50^ models respectively. The diffusion constant computed from the mean squared displacements of a water molecule (shown in supplemental information, SI, Fig. S1) are 5.8× 10^−5^*cm*^2^/*s* for TIP3P and 2.2× 10^−5^*cm*^2^/*s* for TIP4P/2005, respectively. Indeed, the TIP3P estimate exaggerates the bulk water diffusion by a factor of 2.5 while the TIP4P/2005 estimate is nearly identical to the experimental data (2.3× 10^−5^*cm*^2^/*s*). This substantial difference between the two water models naturally leads to the question about the validity of the computation of water conduction through AQPs using the CHARMM force field parameters including the TIP3P parameters. Would TIP4P/2005 give a quantitatively more accurate estimate? The answer should be yes if the water conduction through AQPs is mostly determined by water-water interactions rather than water-protein interactions.

### AQP1/3 permeabilities

In equilibrium, water molecules fill the AQP channels in approximately single files (illustrated in Fig. 2). They constantly fluctuate back and forth inside the channel (on the picosecond time scale) and move out of and into the channel (on the sub-nanosecond time scale). The water flux across the dividing plane (z=0) fluctuates between the positive z- and the negative z-directions to give rise to zero net flux in the absence of an osmotic gradient across the cellular membrane. The single channel permeability *p*_*f*_ quantifies how readily water molecules can move across the membrane through a water channel in response to an osmotic imbalance between the IC and the EC sides. It is equal to the molar volume of water multiplied by the rate of IC-to-EC transition (or, equally, the rate of EC-to-IC transition) given by the TST formula with the correction for recrossing events. We implemented this correction for recrossing events and found the success ratio to be less than 2% for all six sets of simulations (four AQP1/3-RBC-TIP3/4P combinations and two AQP1/3-POPE-TIP3P combinations). The computational details are illustrated in Fig. 3 for the AQP1-RBC-TIP3P model system and in the supplemental information for the other five model systems. The computed values of single channel permeability at 25°C are tabulated in Table III. Two of the six sets are respectively for AQP1 and AQP3 embedded in the model erythrocyte membrane with TIP3P, and similarly another two with the TIP4P/2005 water model. The remaining two sets are respectively for AQP1 and AQP3 embedded in the POPE bilayer with TIP3P. It is interesting to note that the permeability of aquaglyceroporin AQP3 (that conducts glycerol and water) is higher than AQP1 that is water specific and does not conduct glycerol.

**Table III.** Computed AQP1/3 permeabilities *vs.* experimental data on erythrocytes.

|  | In POPE-bilayer,<br>Computed with TIP3P | In RBC membrane,<br>**Computed with TIP3P** | In RBC membrane,<br>Computed with TIP4P |
| --- | --- | --- | --- |
| $p_f^{AQ1}$ per AQP1 monomer | $3.5 \times 10^{-13} \text{ cm}^3/\text{s}$ | **$7.0 \times 10^{-13} \text{ cm}^3/\text{s}$** | $2.4 \times 10^{-13} \text{ cm}^3/\text{s}$ |
| $p_f^{AQ3}$ per AQP3 monomer | $3.7 \times 10^{-13} \text{ cm}^3/\text{s}$ | **$8.1 \times 10^{-13} \text{ cm}^3/\text{s}$** | $4.4 \times 10^{-13} \text{ cm}^3/\text{s}$ |
| Computed RBC permeability* | $(2.1 \pm 0.5^{\#}) \times 10^{-8} \text{ cm}^3/\text{s}$ | **$(4.2 \pm 0.7^{\#}) \times 10^{-8} \text{ cm}^3/\text{s}$** | $(1.7 \pm 0.4^{\#}) \times 10^{-8} \text{ cm}^3/\text{s}$ |
| Experimentally measured RBC permeability ($P_f A$)<sup>§</sup> | **$(4.3 \pm 0.2) \times 10^{-8} \text{ cm}^3/\text{s}$** | | |

\* $P_f A = (N^{AQ1} p_f^{AQ1} + N^{AQ3} p_f^{AQ3})$ with the numbers of AQP1/3 copies per erythrocyte, $N^{AQ1} = (58 \pm 10) \times 10^3$ and $N^{AQ3} = (2.0 \pm 0.5) \times 10^3$ , which were taken from Ref. <sup>40</sup>. <sup>#</sup>The error margins largely reflect those from the proteomics study of Ref. <sup>40</sup> because the simulations gave much smaller error bars.
<sup>§</sup>This experimental value represents the mean between two sets of experiments shown in Fig. 4 (on erythrocytes in 1xPBS) and Fig. 5, top panel (on erythrocytes in 0.7xPBS), respectively.

**Fig. 3.**
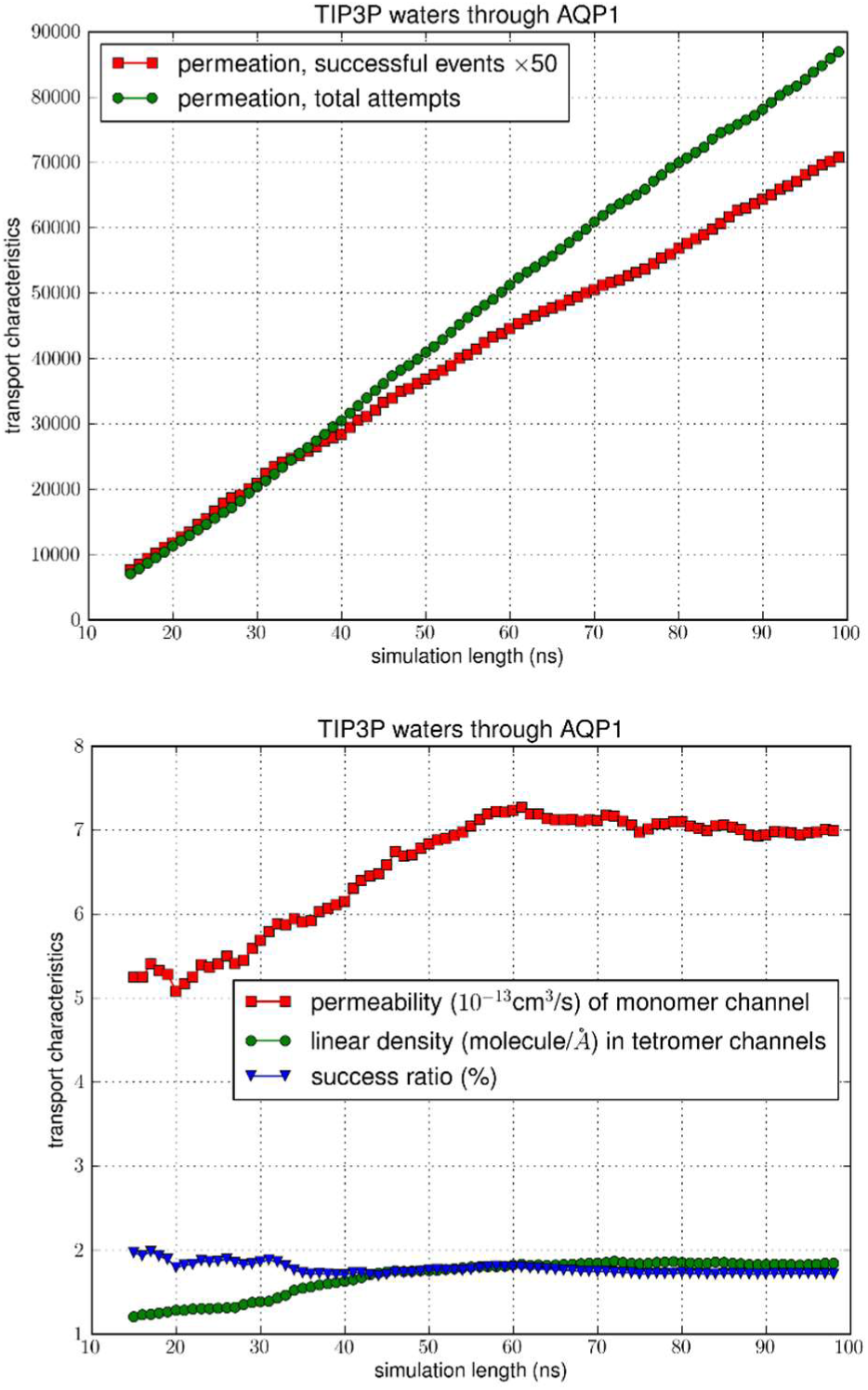
Transport characteristics of erythrocyte AQP1. The top panel shows the time course of the number of transport events *vs.* the number of attempts. The bottom panel shows convergence of the computation: the computed values of permeability, linear density, and success ratio *vs.* length of simulation.

For AQP1/3 in the model erythrocyte membrane, the use of TIP3P and the use of TIP4P/2005 produced results that differ by a factor of about two. These differences are outside the margins of error of our computation and also outside the margin of error of our experimental data. Considering the model erythrocyte membrane *vs.* the lipid bilayer of a single lipid type, the results for AQP1 and AQP3 are also significantly different, outside the margins of error. Therefore, our experiments can be applied to quantitatively validate the AQP1/3-RBC-membrane-TIP3P model.

### Experimental data

To differentiate these *in silico* models, we conducted *in vitro* experiments of erythrocyte shrinking in a hyperosmotic medium probed with the stopped-flow light scattering spectrometer. A small number of (but representative) data points are shown in Fig. 4. From the big data (10,000 data points in each of the 20 or more experimental runs conducted for each condition), we learnt that the process of erythrocyte shrinking in response to hyperosmotic mixing does not precisely follow the exponential behavior (red curve in Fig. 4) that was generally assumed in the current literature. Instead, we can learn the time-course (black curve in Fig. 4) from the data with minimal assumptions that are certainly invariable: the number of AQPs in a given erythrocyte did not change during the shrinkage process and the simple laws of hydrodynamics cannot be altered while the cellular shape and volume are unknown variables. Using the total permeability of a cell *P*_*f*_*A* as the fitting parameter, we obtained the full-dynamics curve along with the mean value and the standard error of *P*_*f*_*A* shown in Fig. 4 and Table III. With this, we can conclude that the *in silico* model system of AQP1/3-RBC-TIP3P is validated by the *in vitro* experiments on human erythrocytes. The other model systems (AQP1/3-POPE-TIP3P or AQP1/3-RBC-TIP4P) deviate from the experimental data by a factor between two and three.

**Fig. 4.**
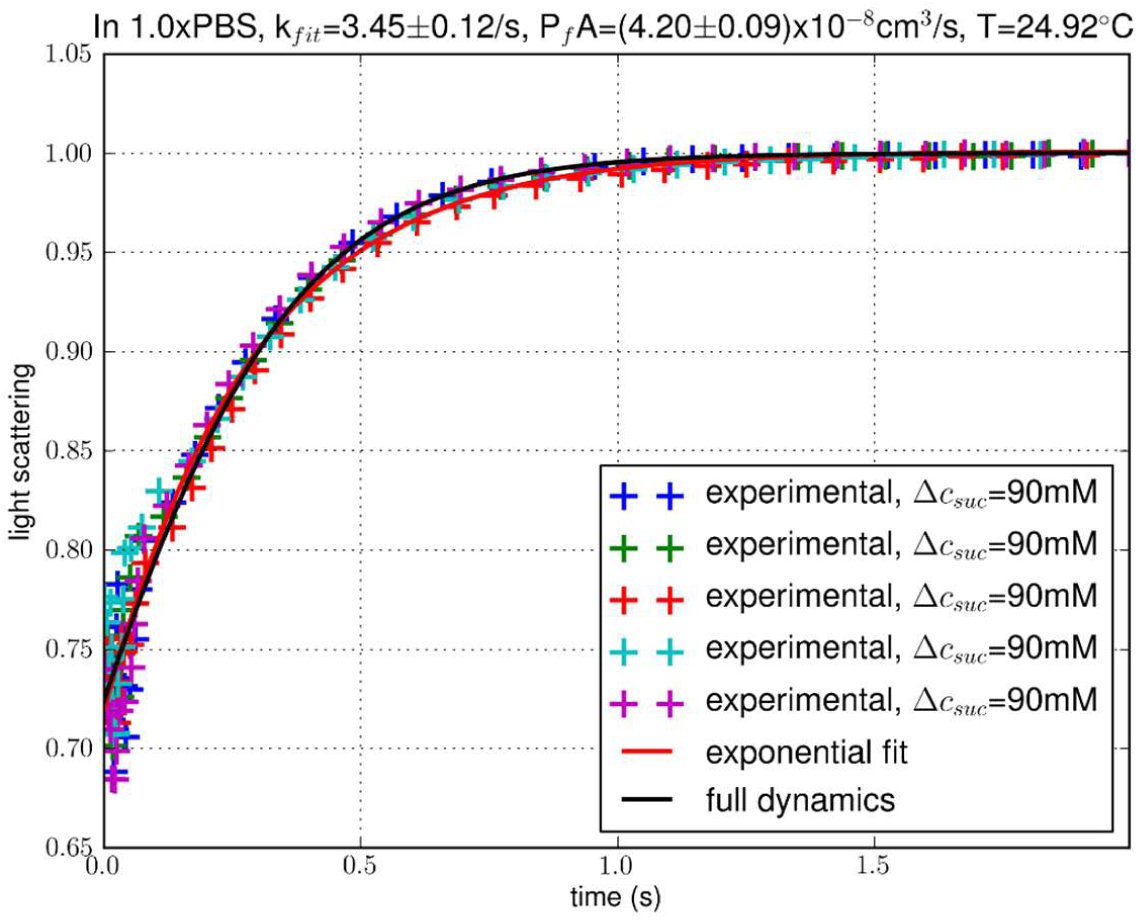
Experimental data of the time courses of human erythrocyte shrinking. The solid curves represent the theoretical predictions from the simple rate equation of osmotic water flux caused by an outward/inward 90 mosM gradient with total water flux *P*_*f*_*A* as a fitting parameter. The crosses represent normalized intensity of 90° scattered light. Light intensity is normalized with the final constant level in each experiment.

It is interesting to examine whether the membrane permeability of erythrocytes depends on the cell volume and conformation. For that purpose, we repeated the erythrocyte shrinking experiments with 0.7× PBS and 1.3× PBS as the buffer liquids respectively. The results shown in Fig. 5, in comparison with Fig. 4, demonstrated that *P*_*f*_*A* is invariant among the three sets of experiments. Considering that the cellular volume and conformation varied significantly from one set of experiments to another, we conclude that water transport across the erythrocyte membrane is, within the margin of error, all through aquaporins. Otherwise, the value of *P*_*f*_*A* would vary with the membrane surface area. We further concluded that aquaporins are not modulated by the cell volume-conformation (illustrated in Fig. 6).

**Fig. 5.**
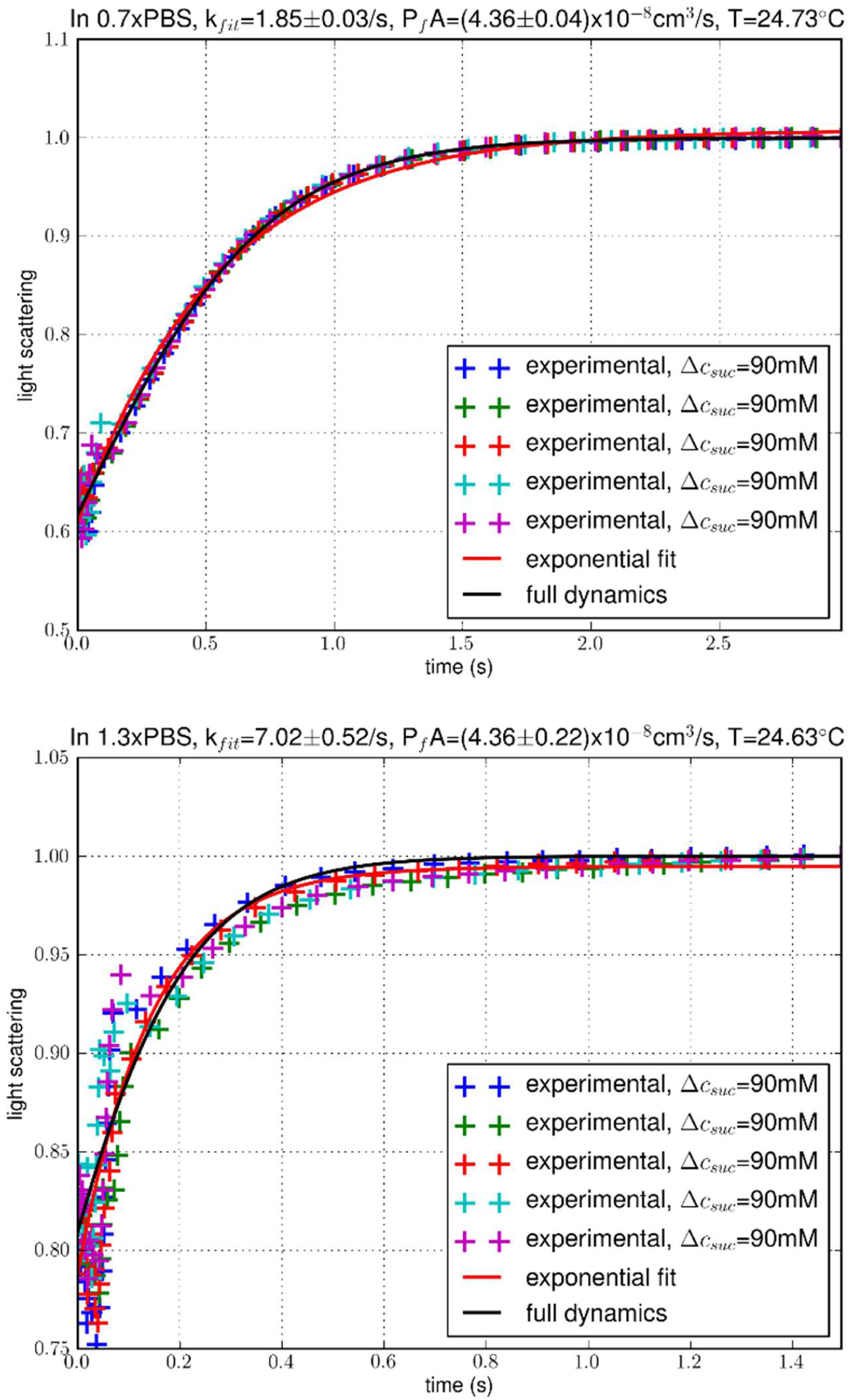
Additional experiments. All conditions are identical to Fig. 4 except that the buffers used were 0.7× PBS (top) and 1.3× PBS (bottom) respectively.

**Fig. 6.**
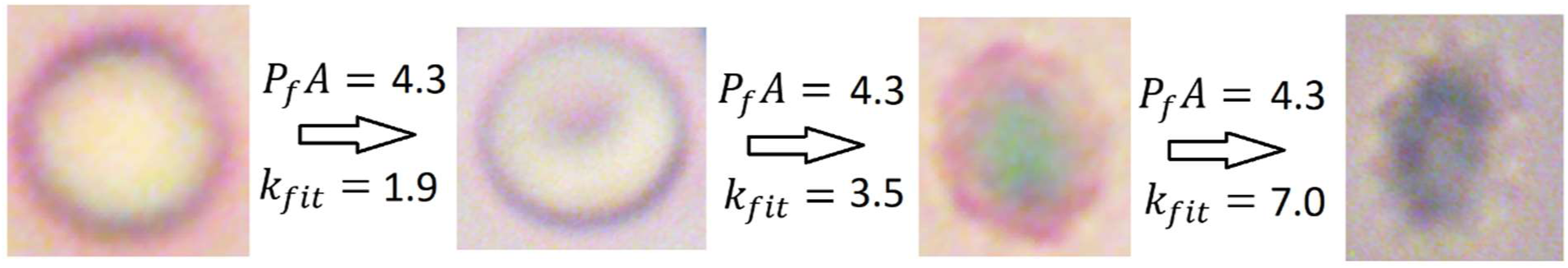
Invariance of AQP permeability illustrated. The numbers indicate total membrane permeability *P*_*f*_*A* = 4.3 × 10^−8^*cm*^3^/*s* and the exponential fitting rate constant *k*_*fit*_ = 1.9/*s*, 3.5/*s, or* 7.0/*s* respectively. The optical images were taken of RBC in 270 mosM (left image), 300 mosM (second from left image), 390 mosM (third from left image), and 480 mosM (right image) saline-sucrose solutions respectively (images not beautified with optical filtering).

### Quantitative significance of lipid compositions

The AQP1/3-RBC-TIP3P simulations produced quantitative agreement with the experimental data while the AQP1/3-POPE-TIP3P simulations underestimated the cellular permeability by a factor of two. Fig. 7 shows how that happened. Looking into the stochastic dynamics of the water molecules interacting with the proteins which are surrounded by the membrane lipids, we noticed that, in a typical frame of the trajectory, all four monomer channels of AQP1 in the model erythrocyte membrane are constantly open for water flux while, on the average, two of the four channels of AQP1 in the POPE bilayer are closed. Why? The erythrocyte membrane has a tighter fit for the membrane proteins, causing AQP1 residues at the ends of the channel to have smaller fluctuations. In contrast, the POPE bilayer is loose, which allows greater fluctuations of the AQP1 residues near the channel ends. This can be seen from the left panels of Fig. 7. Quantitatively, the mean square fluctuations of the channel end residues are shown in SI, Fig. S4. The tight fit of AQP1 in the erythrocyte membrane is in fact optimal to give the greatest permeability of the channel protein.

**Fig. 7.**
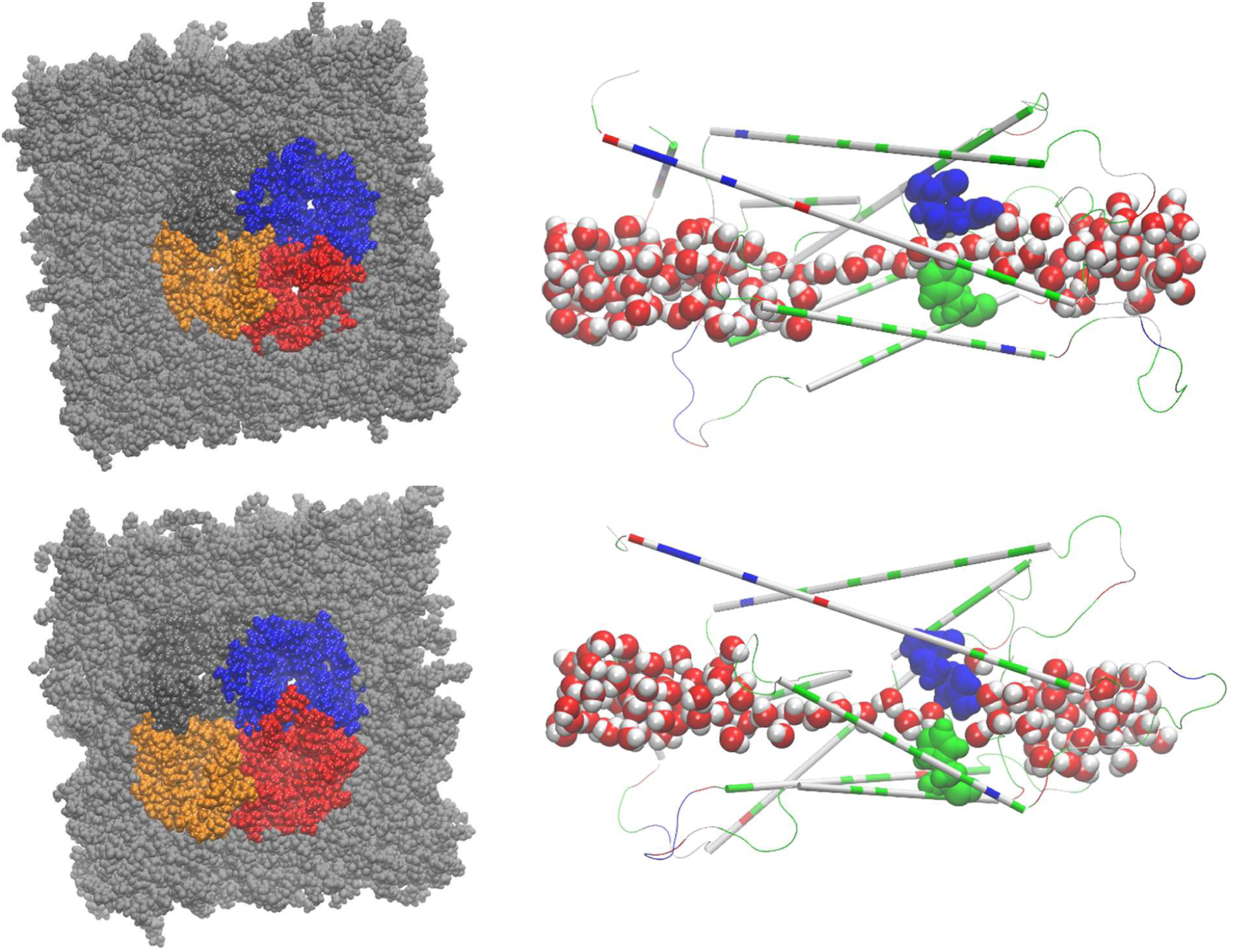
Water conduction through AQP1 in the erythrocyte membrane (top two panels) *vs* that in POPE bilayer (bottom two panels). In the right two panels, the protein monomer is shown in thin cartoons with the ar/R sf residues (Arg 197 and His 182) in large spheres, all colored by residue types (hydrophilic, green; hydrophobic, white; negatively charged, red; positively charged, blue). Water molecules inside and near the channel are shown in large spheres colored by atoms (O, red; H, white). In the left two panels, the protein tetramers (colored by protein monomers) and the membrane lipids (in gray) are all shown in space-filling spheres, viewed from the extracellular side.

## CONCLUSIONS

For water transport through erythrocyte aquaporins, quantitative agreement can be reached between the *in vitro* experiments and the *in silico* simulations if the erythrocyte membrane is modelled with appropriate lipid compositions in its inner and outer leaflets. The TIP3P water model is more accurate than TIP4P for the aquaporin simulations even though the latter gives a better estimate of the bulk water properties. Unlike many other membrane proteins, aquaporins are not modulated by the changes in cellular volume and conformation.

## Supplementary Information

Twelve additional figures and two movies that are discussed but not included in the main text.

## Data availability

The Dataset (parameters, coordinates, scripts, *etc.*) to replicate this study is available at Harvard Dataverse, https://doi.org/10.7910/.

## Author contributions

RC, MF and LYC did the computational work; RC and HL did the experimental measurements; LYC conceptualized the research and wrote the paper; All participated in analyzing the data and editing the manuscript.

## Declaration

There are no conflicts to declare.

## Acknowledgements

The authors acknowledge the computing resources provided by the Texas Advanced Computing Center (TACC) at the University of Texas at Austin.

## REFERENCES

1. G. M. Preston, T. P. Carroll, W. B. Guggino and P. Agre, Science, 1992, 256, 385–387.

2. M. L. Zeidel, S. V. Ambudkar, B. L. Smith and P. Agre, Biochemistry, 1992, 31, 7436–7440.

3. B. Yang and A. S. Verkman, Journal of Biological Chemistry, 1997, 272, 16140–16146.

4. M. Echevarría, E. E. Windhager and G. Frindt, Journal of Biological Chemistry, 1996, 271, 25079–25082.

5. K. Ishibashi, S. Sasaki, K. Fushimi, S. Uchida, M. Kuwahara, H. Saito, T. Furukawa, K. Nakajima, Y. Yamaguchi and T. Gojobori, Proceedings of the National Academy of Sciences, 1994, 91, 6269.

6. N. Roudier, J.-M. Verbavatz, C. Maurel, P. Ripoche and F. Tacnet, Journal of Biological Chemistry, 1998, 273, 8407–8412.

7. T. Zeuthen and D. A. Klaerke, Journal of Biological Chemistry, 1999, 274, 21631–21636.

8. G. Benga, European Biophysics Journal, 2013, 42, 33–46.

9. S. Eriksson, K. Elbing, O. Söderman, K. Lindkvist-Petersson, D. Topgaard and S. Lasič, PLOS ONE, 2017, 12, e0177273.

10. E. Gianolio, G. Ferrauto, E. Di Gregorio and S. Aime, Biochimica et Biophysica Acta (BBA) - Biomembranes, 2016, 1858, 627–631.

11. M. Göllner, A. C. Toma, N. Strelnikova, S. Deshpande and T. Pfohl, Biomicrofluidics, 2016, 10, 054121.

12. M. Palmgren, M. Hernebring, S. Eriksson, K. Elbing, C. Geijer, S. Lasič, P. Dahl, J. S. Hansen, D. Topgaard and K. Lindkvist-Petersson, The Journal of Membrane Biology, 2017, 250, 629–639.

13. M. R. Ruggiero, S. Baroni, S. Pezzana, G. Ferrante, S. Geninatti Crich and S. Aime, Angewandte Chemie International Edition, 2018, 57, 7468–7472.

14. X. Tian, H. Li, X. Jiang, J. Xie, J. C. Gore and J. Xu, Journal of magnetic resonance (San Diego, Calif. : 1997), 2017, 275, 29–37.

15. G. J. Wilson, C. S. Springer, S. Bastawrous and J. H. Maki, Magnetic Resonance in Medicine, 2016, 77, 2015–2027.

16. G. Calamita, J. Perret and C. Delporte, Frontiers in Physiology, 2018, 9, 851.

17. A. Delgado-Bermúdez, M. Llavanera, L. Fernández-Bastit, S. Recuero, Y. Mateo-Otero, S. Bonet, I. Barranco, B. Fernández-Fuertes and M. Yeste, J Anim Sci Biotechnol, 2019, 10, 77–77.

18. A. Delgado-Bermúdez, M. Llavanera, S. Recuero, Y. Mateo-Otero, S. Bonet, I. Barranco, B. Fernandez-Fuertes and M. Yeste, International Journal of Molecular Sciences, 2019, 20.

19. A. Delgado-Bermúdez, F. Noto, S. Bonilla-Correal, E. Garcia-Bonavila, J. Catalán, M. Papas, S. Bonet, J. Miró and M. Yeste, Biology, 2019, 8.

20. K. Gotfryd, A. F. Mósca, J. W. Missel, S. F. Truelsen, K. Wang, M. Spulber, S. Krabbe, C. Hélix-Nielsen, U. Laforenza, G. Soveral, P. A. Pedersen and P. Gourdon, Nature Communications, 2018, 9, 4749.

21. V. Graziani, A. Marrone, N. Re, C. Coletti, J. A. Platts and A. Casini, Chemistry – A European Journal, 2017, 23, 13802–13813.

22. S. Hirako, Y. Wakayama, H. Kim, Y. Iizuka, A. Matsumoto, N. Wada, A. Kimura, M. Okabe, J. Sakagami, M. Suzuki, F. Takenoya and S. Shioda, Obesity Research & Clinical Practice, 2016, 10, 710–718.

23. A. Horner and P. Pohl, Faraday Discussions, 2018, 209, 9–33.

24. U. Laforenza, C. Bottino and G. Gastaldi, Biochimica et Biophysica Acta (BBA) - Biomembranes, 2016, 1858, 1–11.

25. S. W. de Maré, R. Venskutonytė, S. Eltschkner, B. L. de Groot and K. Lindkvist-Petersson, Structure, 2020, DOI: 10.1016/j.str.2019.11.011.

26. M. Arif, P. Kitchen, M. T. Conner, E. J. Hill, D. Nagel, R. M. Bill, S. J. Dunmore, A. L. Armesilla, S. Gross, A. R. Carmichael, A. C. Conner and J. E. Brown, Oncology Letters, 2018, 16, 713–720.

27. D. Posfai, K. Sylvester, A. Reddy, J. G. Ganley, J. Wirth, Q. E. Cullen, T. Dave, N. Kato, S. S. Dave and E. R. Derbyshire, PLoS Pathogens, 2018, 14, e1007057.

28. Z. Zhu, L. Jiao, T. Li, H. Wang, W. Wei and H. Qian, Oncology Letters, 2018, 16, 2661–2667.

29. R. A. Rodriguez, H. Liang, L. Y. Chen, G. Plascencia-Villa and G. Perry, Biochimica et Biophysica Acta (BBA) - Biomembranes, 2019, 1861, 768–775.

30. Y. Sonntag, P. Gena, A. Maggio, T. Singh, I. Artner, M. K. Oklinski, U. Johanson, P. Kjellbom, J. D. Nieland, S. Nielsen, G. Calamita and M. Rützler, Journal of Biological Chemistry, 2019.

31. T. O. Wambo, R. A. Rodriguez and L. Y. Chen, Biochimica et Biophysica Acta (BBA) - Biomembranes, 2017, 1859, 1310–1316.

32. B. L. de Groot and H. Grubmüller, Science, 2001, 294, 2353–2357.

33. F. Zhu, E. Tajkhorshid and K. Schulten, Biophysical Journal, 2004, 86, 50–57.

34. R. J. Law and M. S. P. Sansom, European Biophysics Journal, 2004, 33, 477–489.

35. M. Hashido, M. Ikeguchi and A. Kidera, FEBS Lett., 2005, 579, 5549–5552.

36. M. Ø. Jensen and O. G. Mouritsen, Biophys. J., 2006, 90, 2270–2284.

37. M. Hashido, A. Kidera and M. Ikeguchi, Biophys. J., 2007, 93, 373–385.

38. J. S. Hub and B. L. de Groot, Proc. Natl. Acad. Sci. U.S.A., 2008, 105, 1198–1203.

39. J. A. Freites, K. L. Németh-Cahalan, J. E. Hall and D. J. Tobias, Biochimica et Biophysica Acta (BBA) - Biomembranes, 2019, 1861, 988–996.

40. A. H. Bryk and J. R. Wiśniewski, Journal of Proteome Research, 2017, 16, 2752–2761.

41. J. A. F. Op den Kamp, Annual Review of Biochemistry, 1979, 48, 47–71.

42. A. A. Spector and M. A. Yorek, Journal of Lipid Research, 1985, 26, 1015–1035.

43. J. A. Virtanen, K. H. Cheng and P. Somerharju, Proceedings of the National Academy of Sciences, 1998, 95, 4964.

44. M. Aoun, P. A. Corsetto, G. Nugue, G. Montorfano, E. Ciusani, D. Crouzier, P. Hogarth, A. Gregory, S. Hayflick, G. Zorzi, A. M. Rizzo and V. Tiranti, Molecular Genetics and Metabolism, 2017, 121, 180–189.

45. S. Himbert, R. J. Alsop, M. Rose, L. Hertz, A. Dhaliwal, J. M. Moran-Mirabal, C. P. Verschoor, D. M. E. Bowdish, L. Kaestner, C. Wagner and M. C. Rheinstädter, Scientific Reports, 2017, 7, 39661.

46. L. Y. Chen and C. F. Phelix, Biochemical and Biophysical Research Communications, 2019, 511, 573–578.

47. W. L. Jorgensen, J. Chandrasekhar, J. D. Madura, R. W. Impey and M. L. Klein, The Journal of Chemical Physics, 1983, 79, 926–935.

48. E. E. S. Ong and J.-L. Liow, Fluid Phase Equilibria, 2019, 481, 55–65.

49. A. Zaragoza, M. A. Gonzalez, L. Joly, I. López-Montero, M. A. Canales, A. L. Benavides and C. Valeriani, Physical Chemistry Chemical Physics, 2019, 21, 13653–13667.

50. J. L. F. Abascal and C. Vega, The Journal of Chemical Physics, 2005, 123, 234505.

51. W. Humphrey, A. Dalke and K. Schulten, Journal of Molecular Graphics, 1996, 14, 33–38.

52. S. Jo, T. Kim, V. G. Iyer and W. Im, J. Comput. Chem., 2008, 29, 1859–1865.

53. J. C. Phillips, R. Braun, W. Wang, J. Gumbart, E. Tajkhorshid, E. Villa, C. Chipot, R. D. Skeel, L. Kalé and K. Schulten, Journal of Computational Chemistry, 2005, 26, 1781–1802.

54. R. B. Best, X. Zhu, J. Shim, P. E. M. Lopes, J. Mittal, M. Feig and A. D. MacKerell, Journal of Chemical Theory and Computation, 2012, 8, 3257–3273.

55. K. Vanommeslaeghe, E. Hatcher, C. Acharya, S. Kundu, S. Zhong, J. Shim, E. Darian, O. Guvench, P. Lopes, I. Vorobyov and A. D. Mackerell, Journal of Computational Chemistry, 2010, 31, 671–690.

56. J. B. Klauda, R. M. Venable, J. A. Freites, J. W. O’Connor, D. J. Tobias, C. Mondragon-Ramirez, I. Vorobyov, A. D. MacKerell and R. W. Pastor, The Journal of Physical Chemistry B, 2010, 114, 7830–7843.

57. X. Zhou and F. Zhu, Journal of Chemical Information and Modeling, 2019, 59, 777–785.

58. N. E. Henriksen and F. Y. Hansen, Physical Chemistry Chemical Physics, 2002, 4, 5995–6000.

59. M. Harri, A.-N. Tapio and J. Hannes, Efficient dynamical correction of the transition state theory rate estimate for a flat energy barrier, 2018.

60. D. Chandler, The Journal of Chemical Physics, 1978, 68, 2959–2970.

61. A. Horner, F. Zocher, J. Preiner, N. Ollinger, C. Siligan, S. A. Akimov and P. Pohl, Science Advances, 2015, 1, e1400083.

