## Supplemental figures for "*In silico* simulations of erythrocyte aquaporins with quantitative *in vitro* validation"

Here we present additional figures that are referred to but not included in the main text:

In Fig. S1, we show the simulation data of bulk water diffusion with TIP3P vs TIP4P.

In Figs. S2a and b, we show the simulation data of AQP1 embedded in POPE bilayer;

In Figs. S2c and d, we show the simulation data of erythrocyte AQP1 but water modelled with TIP4P;

In Figs. S3a and b, we show the simulation data of erythrocyte AQP3;

In Figs. S3c and d, we show the simulation data of AQP3 embedded in POPE bilayer;

In Figs. S3e and f, we show the simulation data of erythrocyte AQP3 but water modelled with TIP4P;

In Fig. S4, we show fluctuations of the non-helix residues of AQP1 in erythrocyte membrane *vs* in POPE bilayer.

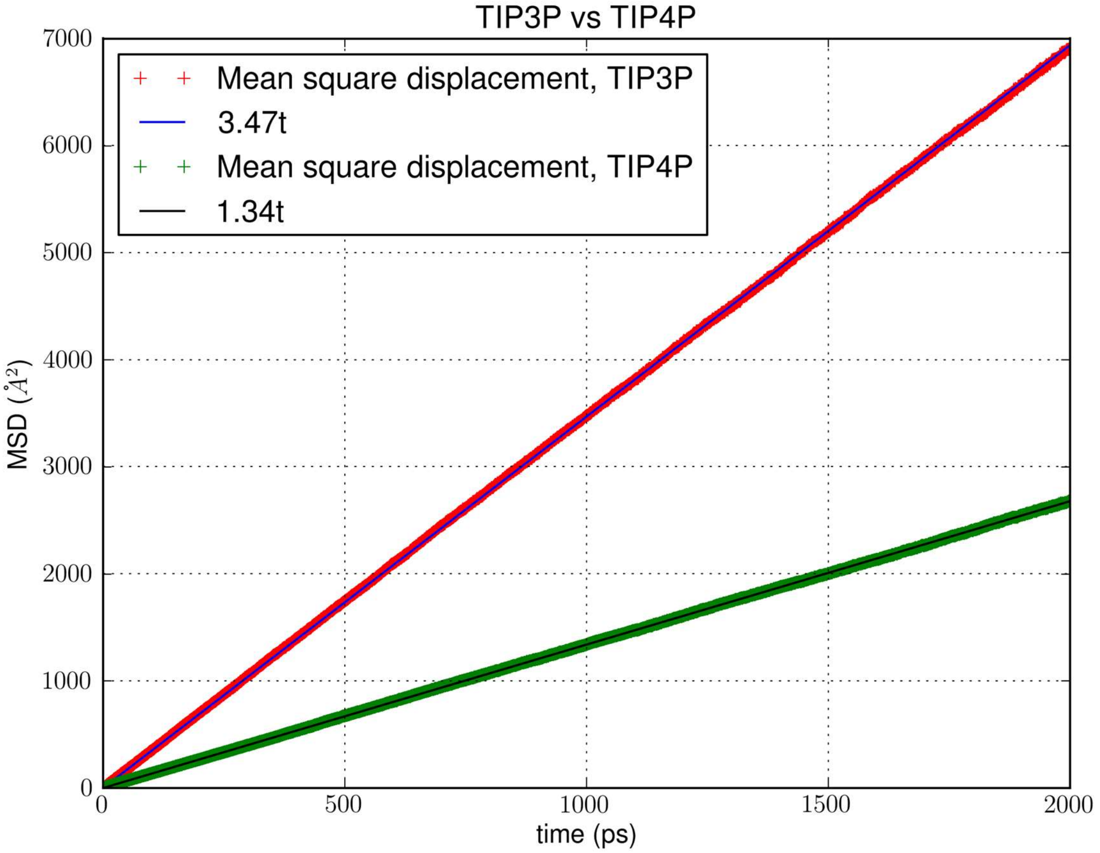


**Fig. S1.** Diffusion in bulk water modelled as TIP3P *vs* TIP4P: Mean squared displacement of a water molecule as a function of the simulation time. For each water model, the simulation was 100 ns in length and was folded into 50 sets of data over 2000 ps.

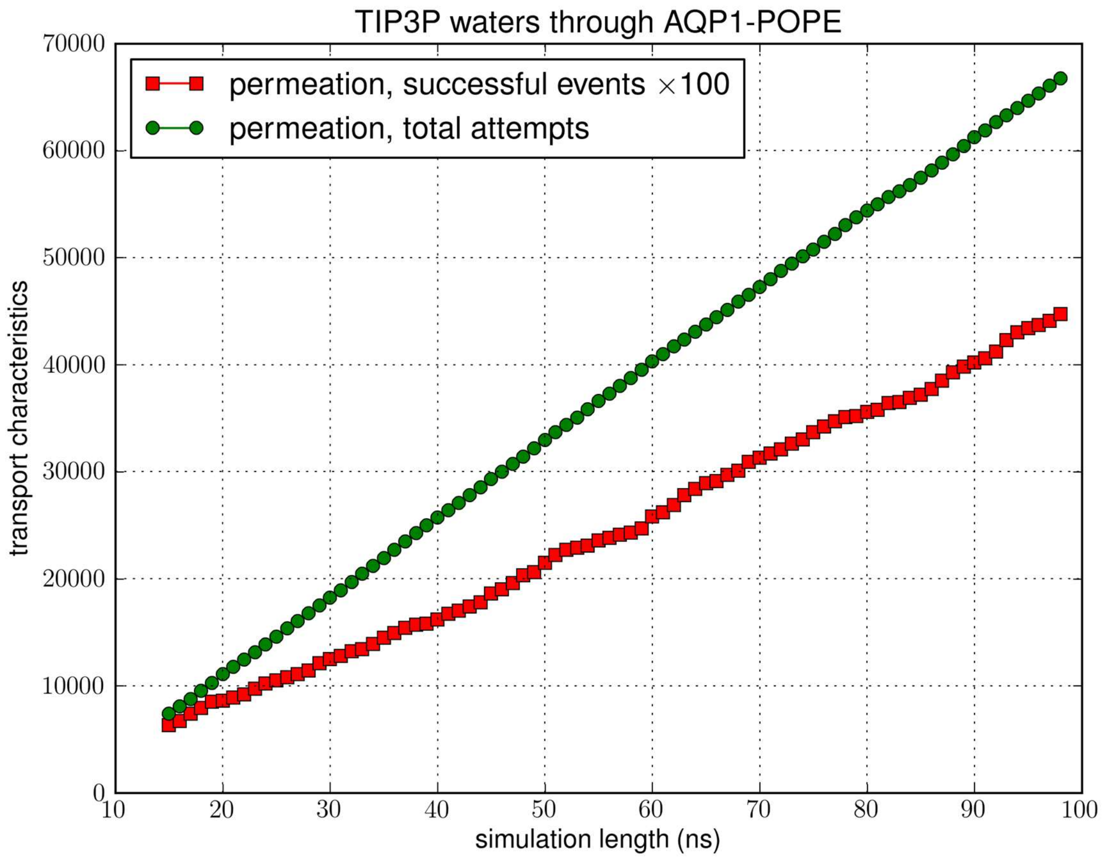


**Fig. S2a**. Transport characteristics of AQP1 embedded in POPE bilayer: the time course of the number of transport events *vs.* the number of attempts.

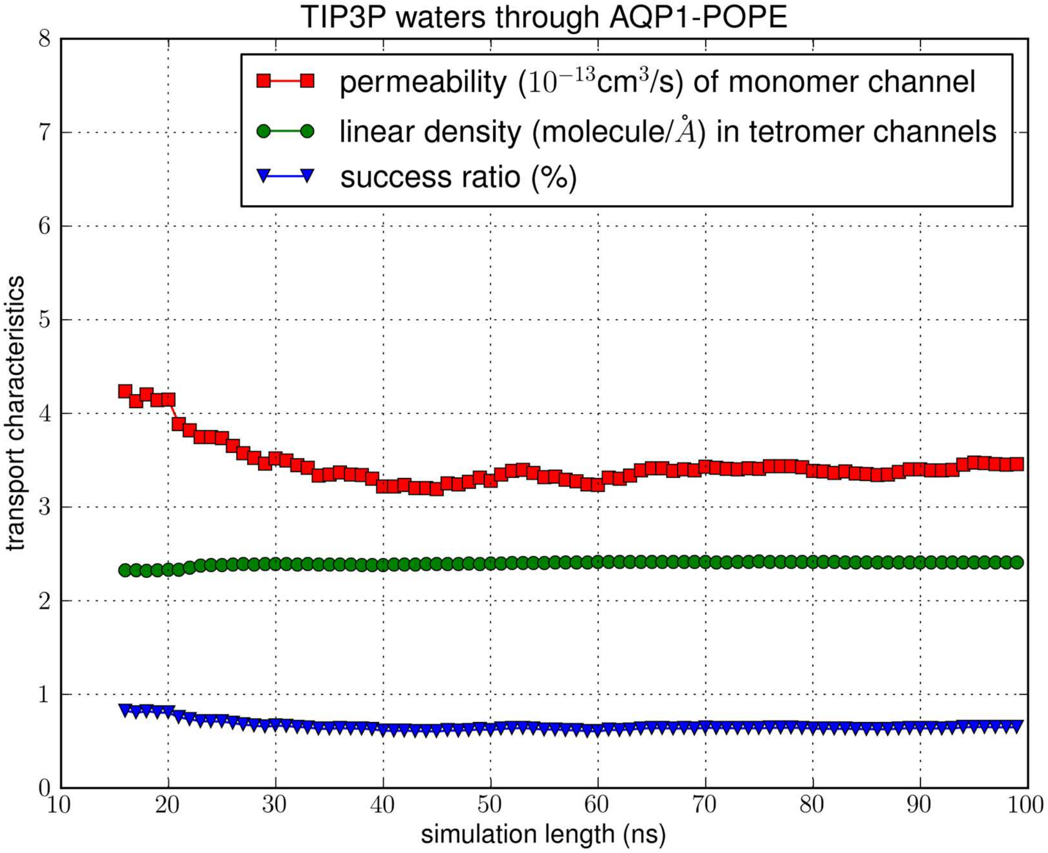


**Fig. S2b**. Transport characteristics of AQP1 embedded in POPE bilayer: convergence of the computation: the computed values of permeability, linear density, and success ratio *vs.* length of simulation.

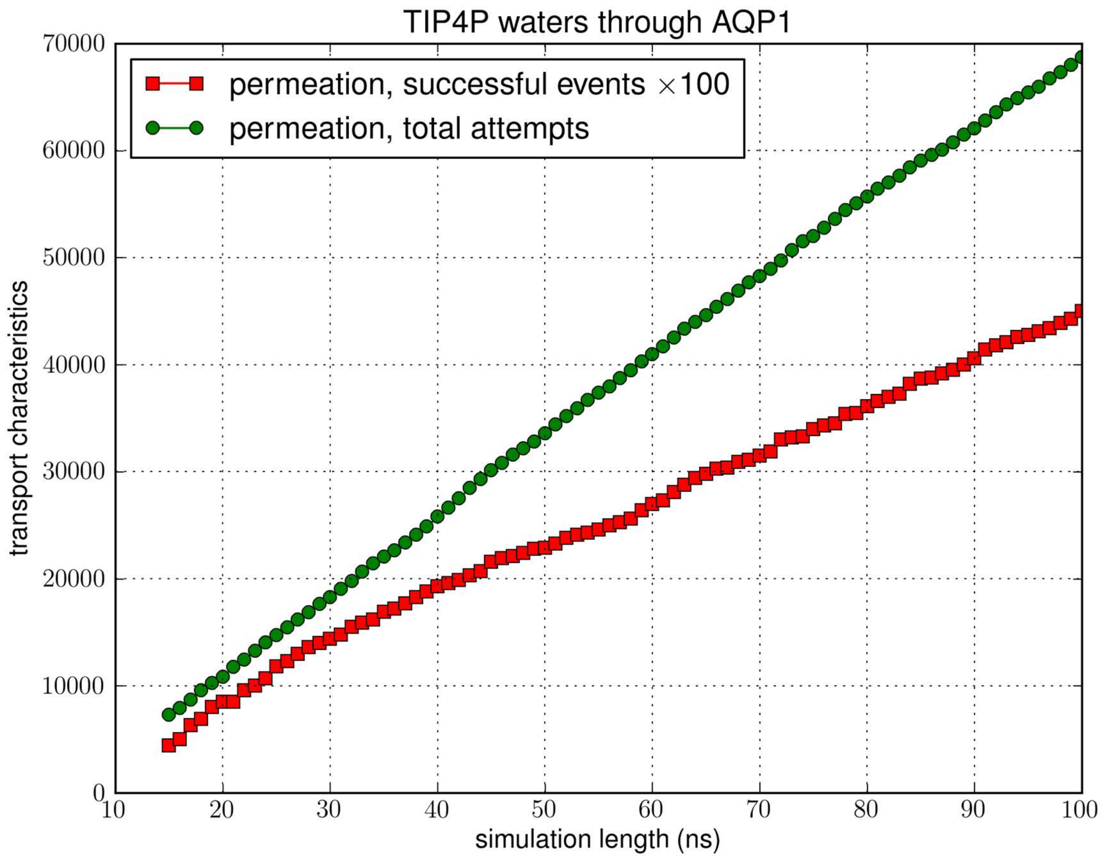


**Fig. S2c**. Transport characteristics of erythrocyte AQP1 with TIP4P water model: the time course of the number of transport events *vs.* the number of attempts.

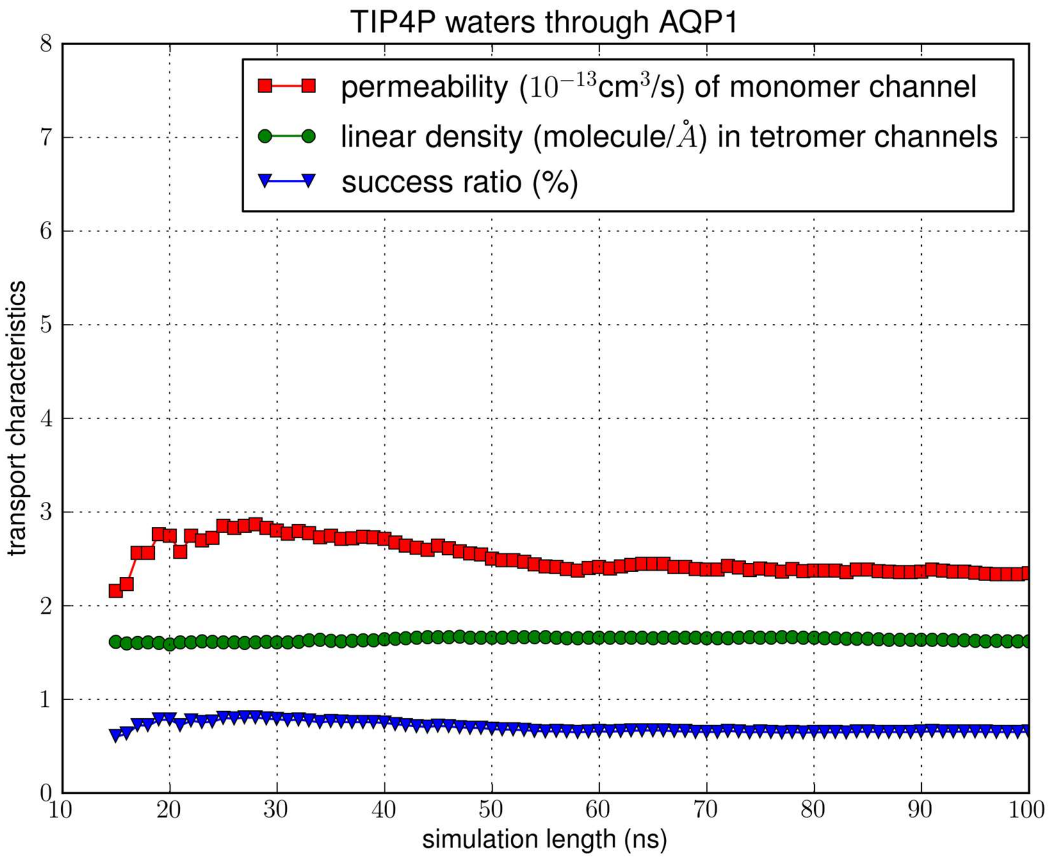


**Fig. S2d**. Transport characteristics of erythrocyte AQP1 with TIP4P water model: convergence of the computation: the computed values of permeability, linear density, and success ratio *vs.* length of simulation.

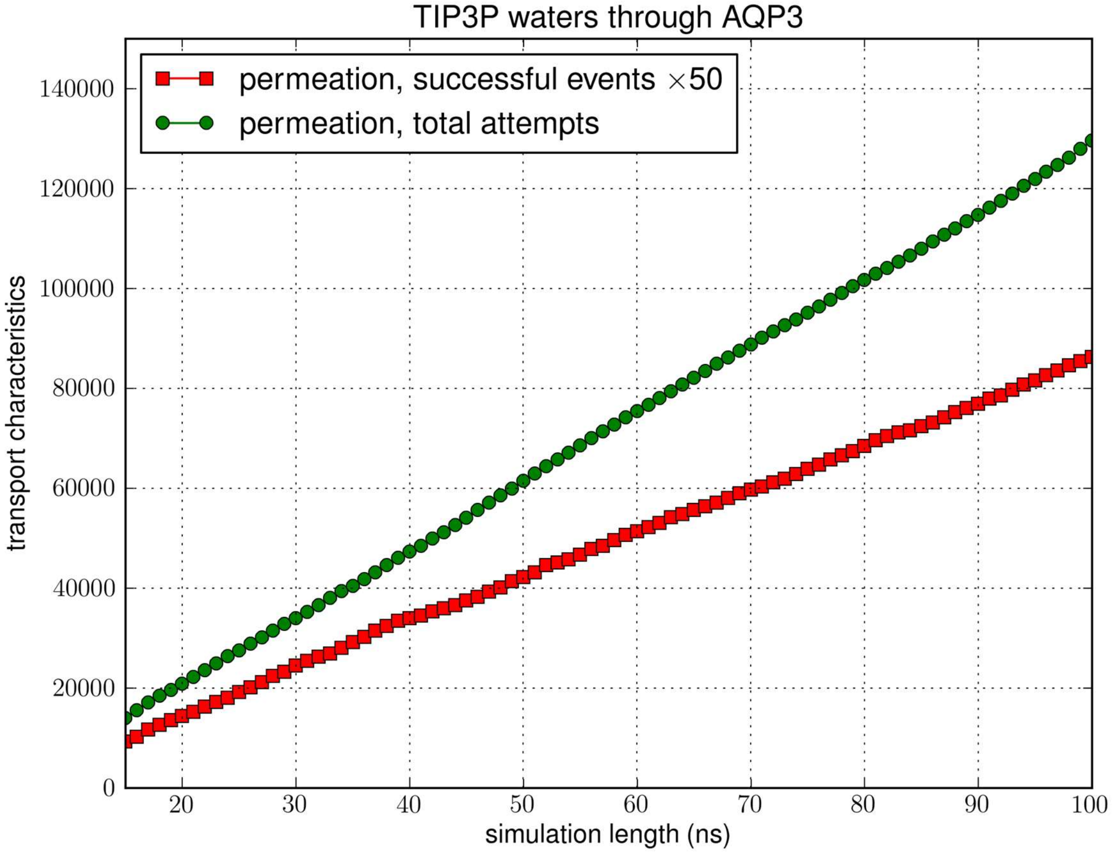


**Fig. S3a**. Transport characteristics of erythrocyte AQP3: the time course of the number of transport events *vs.* the number of attempts.

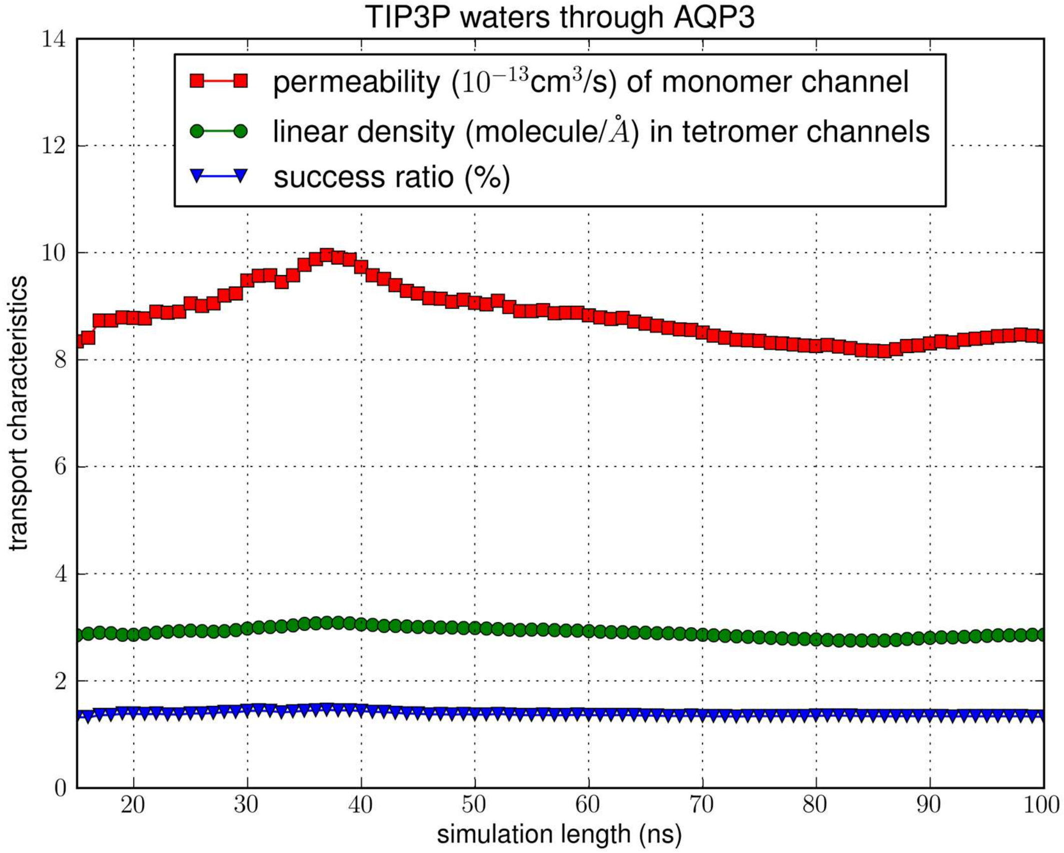


**Fig. S3b**. Transport characteristics of erythrocyte AQP3: convergence of the computation: the computed values of permeability, linear density, and success ratio *vs.* length of simulation.

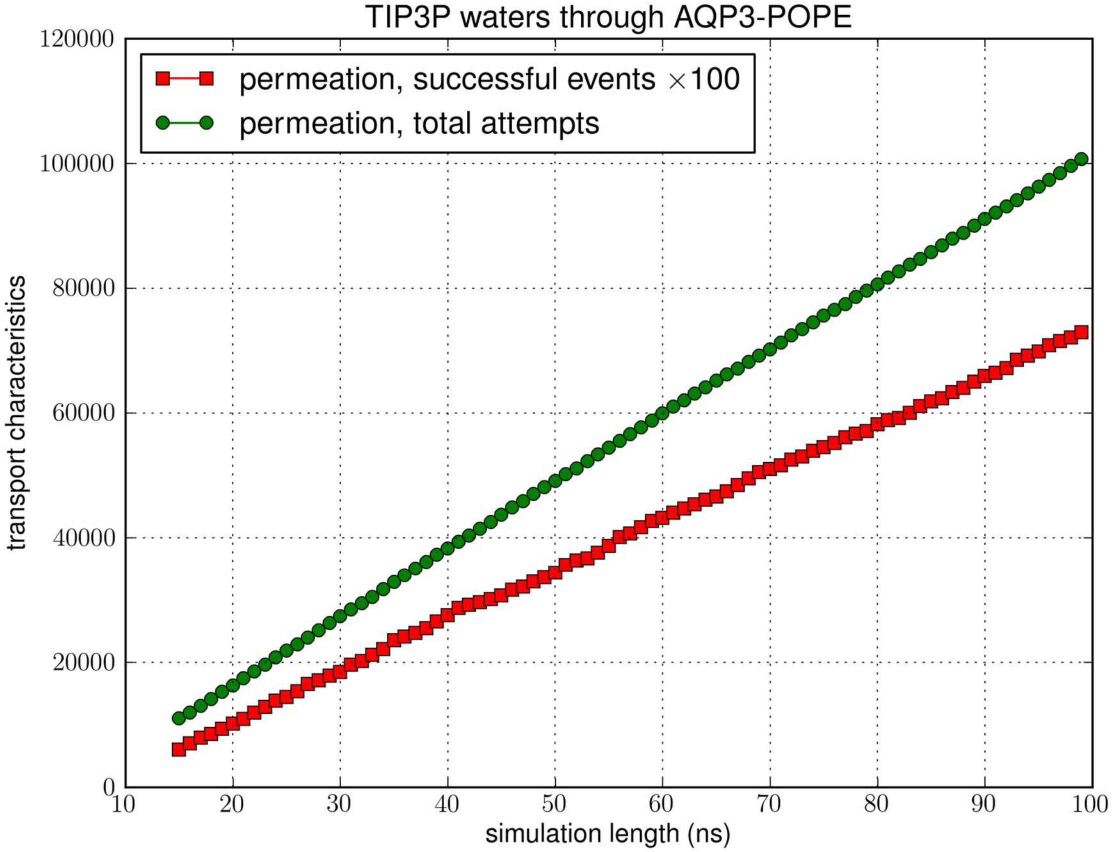


**Fig. S3c**. Transport characteristics of AQP3 embedded in POPE bilayer: the time course of the number of transport events *vs.* the number of attempts.

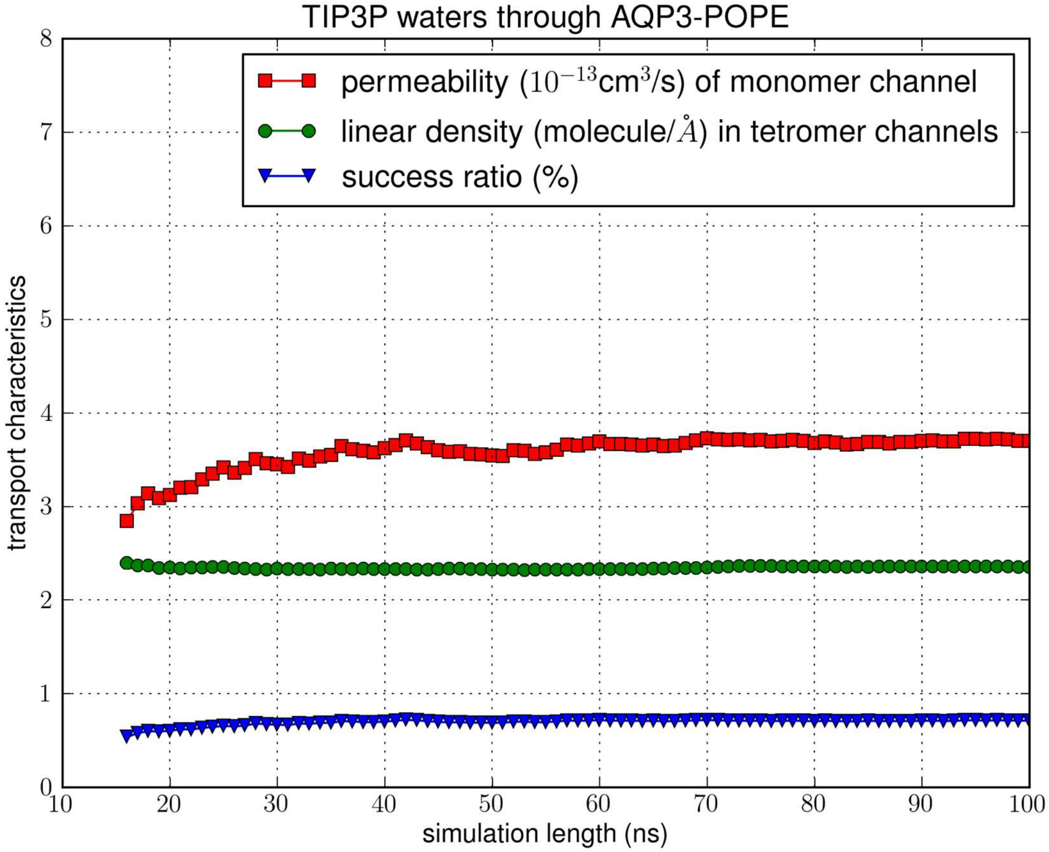


**Fig. S3d**. Transport characteristics of AQP3 embedded in POPE bilayer: convergence of the computation: the computed values of permeability, linear density, and success ratio *vs.* length of simulation.

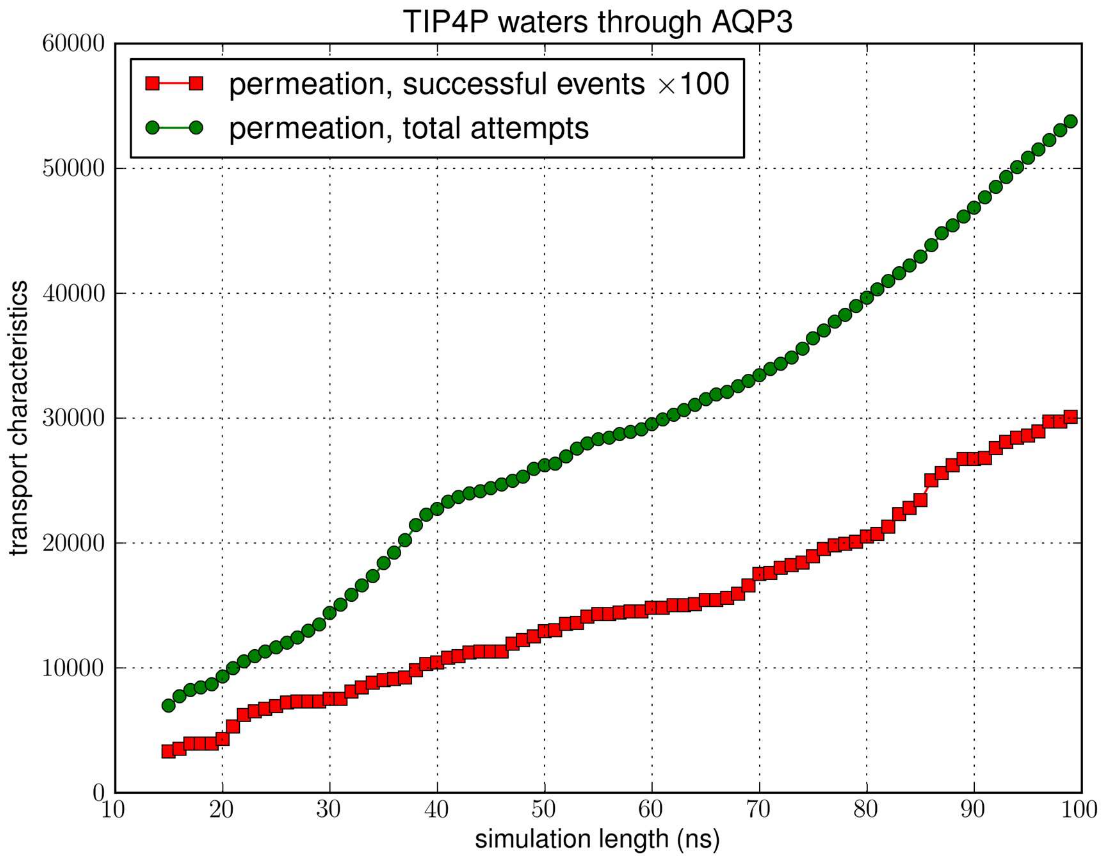


**Fig. S3e**. Transport characteristics of erythrocyte AQP3 with TIP4P water model: the time course of the number of transport events *vs.* the number of attempts.

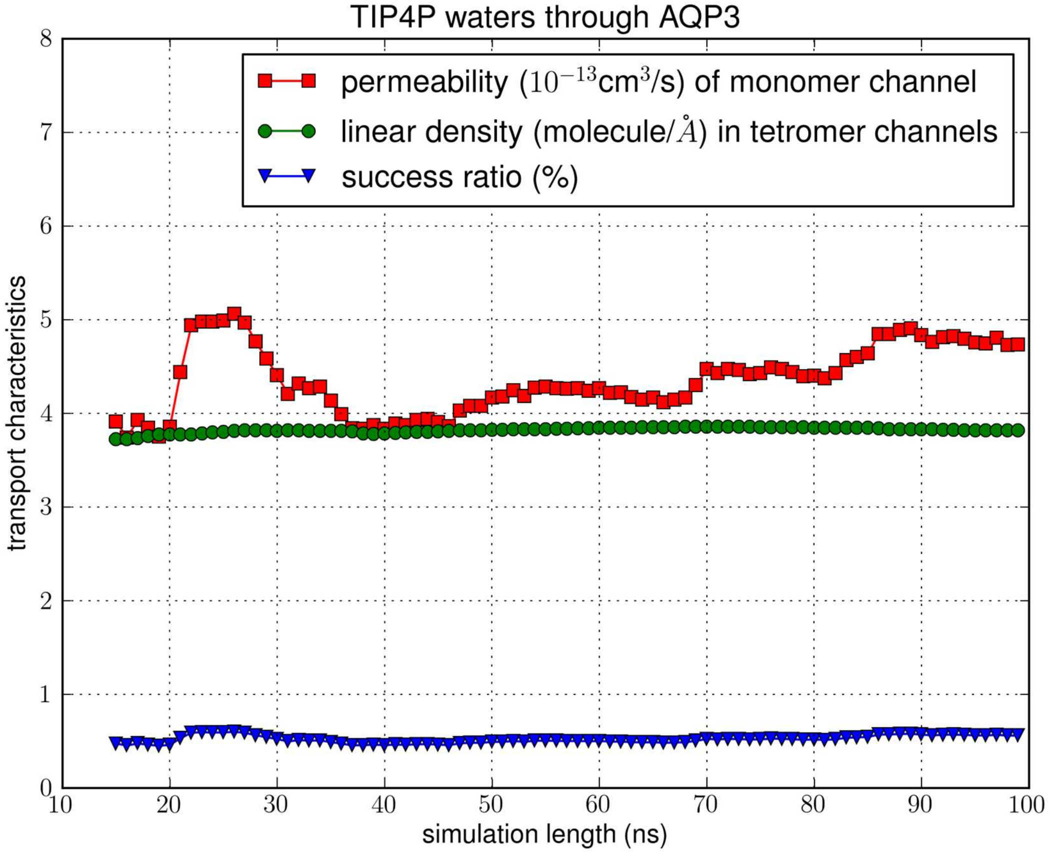


**Fig. S3f**. Transport characteristics of erythrocyte AQP3 with TIP4P water model: convergence of the computation: the computed values of permeability, linear density, and success ratio *vs.* length of simulation.

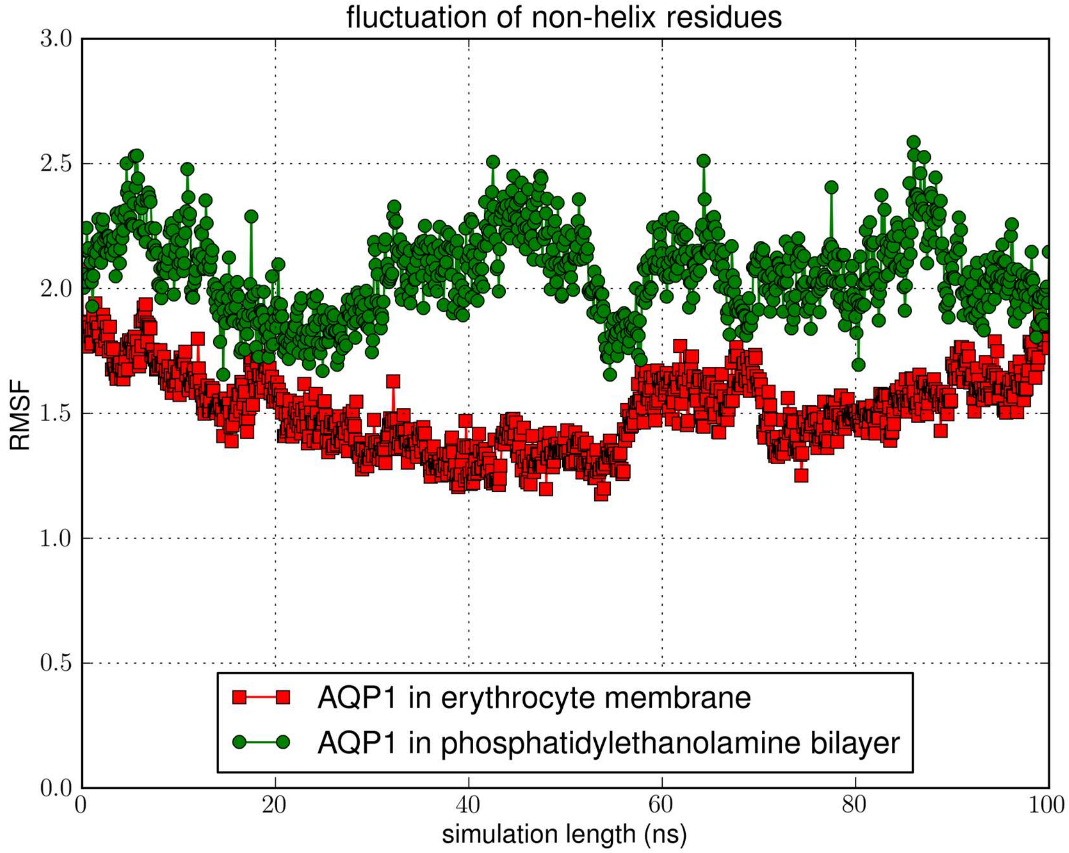


**Fig. S4**. Fluctuations of the non-helix residues of AQP1 in erythrocyte membrane *vs* in POPE bilayer.
